# Flexibility in task performance and division of labour in a cooperatively breeding cichlid

**DOI:** 10.1101/2025.09.08.674851

**Authors:** A. Ramesh, B. Taborsky

## Abstract

Division of labour is a key feature of cooperative social systems, where task specialization among individuals enhances group efficiency. In the cooperatively breeding cichlid *Neolamprologus pulcher* reproductive division of labour exists, where a dominant breeding pair reproduces while subordinates help in rearing the offspring and perform various tasks to gain acceptance within the group. Larger helpers engage in territory maintenance and predator defence while smaller helpers focus on egg care and deterrence of egg predators. Here we investigated task specialisation, division of labour and the dynamics of coordination of tasks in *N. pulcher* groups of natural size and composition. In lab experiments, we assessed whether helpers consistently specialized in sand removal from territories or in egg predator defence when both tasks were presented simultaneously. While different size classes performed both tasks, task performance was not repeatable, and there was no clear division of labour. Dominant females did most work, with the helpers often remaining idle. Lag sequence analysis revealed that individuals were significantly more likely to take up a task if it had just been performed by another individual, rather than dividing labour between individuals - a phenomenon we term “task contagion”. This suggests that individuals respond to immediate group needs, offering new insights into how cooperative breeders can adapt to changing task demands by flexible behaviour and potentially enhance group efficiency.

**Significance statement:** Cooperatively breeding vertebrates exhibit complex social structures and group dynamics, one notable feature being division of labour. However, unlike eusocial insects, these vertebrates display a high degree of flexibility in task allocation among group members. In a laboratory experiment, we investigated the dynamics of task allocation and coordination in groups of the cooperatively breeding cichlid *Neolamprologus pulcher*. We simultaneous induced demands for both territory maintenance and territory defence against egg predators. Task performance and the degree of specialisation varied across size classes and between different groups, and were not repeatable. Using high-resolution event data, we identified a tendency for individuals to follow others in the execution of the same task, a phenomenon we have termed ‘task contagion’.

## Introduction

Sociality is ubiquitous among many animal taxa, which organize into groups of varying sizes and complexities (Frank, 2007). Living in groups almost always leads to emergent properties, even under experimentally created conditions when all animals appear to be identical, due to non-additive or synergistic interactions among individuals (Viscido *et al*., 2004; Wood & Ackland, 2007; Kolokolnikov *et al*., 2013). One of the most striking and complex emergent property of social groups is division of labour (DoL), wherein different group members perform different tasks or roles either permanently, temporarily or spontaneously (Taborsky 2025)s. Division of labour – broadly defined as an uneven distribution of tasks among cooperating individuals – is known for its pivotal role in the success of social organisms and considered one of the key steps in major evolutionary transitions (Szathmáry & Smith, 1995).

Classical examples of division of labour were described in eusocial insects, which can exhibit both reproductive division of labour between reproductive castes and workers as well as non-reproductive division of labour among the workers. In some cases, workers specialize into morphologically distinct castes (such as worker and soldier termites, Engel *et al*., 2016). In the absence of morphological specialization, variation in task performance can correlate with age (such as in honeybee workers, which progress from brood care to foraging task as they age, Seeley, 1982), size or weight, and can be manifested by variation in response thresholds (Rittschof & Robinson, 2013; Jeanson & Weidenmüller, 2014; Beshers & Fewell, 2025). DoL is generally thought to confer efficiency and fitness benefits to the group by individual specialisation, although this depends on the effective coordination among group members (Oster & Wilson, 1979). While specialization is assumed to improve individual skill, the necessity to change tasks can also impose switching costs (Chittka & Muller, 2009; Jeanson & Lachaud, 2015). Rigid task specialisation can be disadvantageous in the case of environmental disturbance or a change in demand for work within a group for other reasons. Thus, a balance between specialisation and flexibility is essential. Indeed, empirical and theoretical work suggests that a compromise between specialisation and behavioural plasticity is necessary to maintain group performance and that in social species, individuals typically maintain some level of flexibility (Dornhaus, 2008; Jeanson, 2019).

Although most work on DoL has been done in eusocial insects, task specialisation and labour division also occurs in some cooperatively-breeding vertebrates. In these species, task allocation and division of labour exhibits significant flexibility, with variation in overall workload among individuals, allowing them to engage in multiple tasks to varying extents (Smith & Riehl, 2022). Instead of strict specialisation or morphological castes, task allocation and social roles are more flexibly determined in cooperative vertebrates and depend on body size, age, experience, and social status (Heinsohn & Legge, 1999; Arnold *et al*., 2005). For example, there is helper specialisation in the social roles of meerkats, *Suricata suricatta*, into sentinels or caregivers (Clutton-Brock *et al*., 2004), and provisioning vs predator deterrence in the superb fairy-wrens, *Malurus cyaneus*, and grey-crowned babblers, *Pomatostomus temporalis* (Russell 2004; Gardner et al. 2003); while in Damaraland mole rats, *Fukomys damarensi*, older individuals handle more demanding tasks compared to younger ones (Zöttl *et al*., 2016; Thorley *et al*., 2018). In the cichlid *Neolamprologus pulcher*, helper contributions are influenced by social dynamics and hierarchy (Naef & Taborsky, 2020a; b; García-Ruiz & Taborsky, 2024), all of which highlight how spatial distribution impacts flexible division of labour (Bruintjes & Taborsky, 2011).

When tasks are assigned within a group, DoL can manifest in a multitude of ways and irrespective of individual specialisation (for example, due to resource sharing or social dynamics among identical or generalist individuals (Kreider *et al*., 2022; Tate Holbrook *et al*., 2009). Individuals may appear to execute tasks in succession, suggesting the presence of a priority system based on urgency or importance of tasks. Tasks may also need to be performed by multiple individuals together to be effective, such as mobbing or defence against predators (Graw & Manser, 2007; Jungwirth *et al*., 2015a; Engelhardt *et al*., 2025). Interestingly, there can also be instances of spontaneous division of labour, wherein individuals distribute amongst tasks over very short time instances without overall specialisation by real-time behavioural adjustments (Taborsky, 2025b). Further, there may be a ‘reserve’ set of individuals who appear idle but can quickly step in when demands for a task suddenly rise (Charbonneau & Dornhaus, 2015; Leitner & Dornhaus, 2019). This highlights that there can be flexible responses to task demands, involving both the availability of workforce and the coordination among individuals. This is crucial in systems with small group sizes, which experience rapid shifts in demands and workforce because they are less well buffered against environmental changes (Netz *et al*., 2025). Despite evidence for both flexibility and division of labour, how cooperative breeders coordinate task performance within groups remains largely unexplored.

In the cooperatively breeding cichlids, *N. pulcher*, groups typically consists of a dominant breeding pair, comprised of the largest individuals, and 1-25 related or unrelated subordinate helpers, organised in a size-based hierarchy. Group members face different task demands including territory defence against conspecific intruders, predator deterrence, brood care, territory maintenance. Although in principle, all individuals in a group are able to perform all these tasks, previous field studies have shown that that body size, the location where a task is encountered, and dominance status of helpers influences the task allocation and division of labour in these cichlids. Small helpers tended to perform brood defence close to the breeding cavity, while large helpers and female breeders devoted more effort to digging out the shelter and engaging in defence against larger predators further from the breeding cavity (Bruintjes & M. Taborsky, 2011; Heg & M. Taborsky, 2010). In addition, we expect work load to be unevenly distributed among the breeders and the different helper sizes – breeder females are known to have the highest workload (Balshine *et al*., 2001). Further, increasing group sizes can reduce the workload on the breeders (Balshine *et al*., 2001; Josi *et al*., 2020) and the relative investment in the helper workload depends on their relatedness to the male or female breeder and their behaviour types and submission rates (Bergmüller & Taborsky, 2005; Stiver *et al*., 2005; Le Vin *et al*., 2011; Zöttl *et al*., 2013). Moreover, unrelated individuals show task-coordination and reciprocate prior helping experience when they need to cooperatively dig out a common shelter, especially after predator exposure (Taborsky & Riebli, 2020). This makes this species a highly suitable system to investigate how work is distributed and organized among group members.

In a controlled lab experiment, we aimed to verify previous findings from the field regarding size-based task allocation and DoL. Further, by collecting fine-resolution event data, we explored patterns of task coordination over short and longer time scales. To achieve this, we created eleven groups consisting of one breeding pair and eight helpers of three different size classes. We investigated helping behaviour across size classes when groups are faced with two simultaneous tasks, digging sand away from shelters and defending against the egg predator *Telmatochromis vittatus*, which was repeated three times. We then computed the task specialisation per size-class (using Proportional Similarity Index) and the consistency of task allocation among size classes within groups and DoL (using Normalised Mutual Entropy) across repeats. To investigate task coordination within trials, we used the fine-resolution behavioural data to perform a lag-sequence analysis, which allows us to create a network of task transitions across different size classes. Here we examined: (a) the likelihood of individuals/size classes to switch tasks within a trial and the direction of these transitions (digging to defense or defense to digging), and (b) whether patterns emerged consistent with spontaneous division of labour among group members, where a helper is more likely to perform task A when another is occupied with task B, or with behavioural contagion, where the likelihood of an individual performing task A increases when another one is already engaged in it.

## Methods

### Study species and housing

*N. pulcher* is a cooperatively breeding cichlid endemic to Lake Tanganyika, East Africa (Konings, 1998). A typical group consists of a dominant breeding pair, comprised of the largest individuals, and 1-25 related or unrelated subordinate helpers. Groups are organized into linear, size-based hierarchies (Dey *et al*., 2013). Helpers ‘pay rent’ to be allowed to stay in the territory by engaging in cooperative tasks in the group such as territory defence against predators, territory maintenance and alloparental care of eggs and larvae (Taborsky & Limberger, 1981). Upon maturation, subordinates face a decision to disperse to neighbouring groups, where they can join at a better rank position or take over a dominant position or to remain in their natal territory to potentially inherit the territory and breeding position in the future (Stiver *et al*., 2007; Jungwirth *et al*., 2015b, 2023; Fischer *et al*., 2017)

### Housing conditions

The experiments were conducted at the Ethologische Station Hasli of the Institute for Ecology and Evolution, University of Bern, Switzerland. The experimental fish were taken from a laboratory-bred stock originally derived from wild-caught fish from the southern end of Lake Tanganyika near Mpulungu, Zambia. Individuals used for the experiment were taken from stock tanks containing 20 – 50 fish, housed in 200L - 400L tanks (Reyes-Contreras et al. 2023). Fish were housed under conditions mimicking their natural habitat with respect to the biochemical parameters, light conditions (L:D 13:11 h, with 10 min of dimmed light in the mornings and evenings) and the water temperature (27 ± 1°C). The fish were fed once a day with commercial food flakes (5 days a week) and frozen zooplankton (1 day a week). TetraMin Baby® powdered flakes were added when fry were present.

### Experimental groups

For the experiment, eleven social groups, composed of ten individuals per group, were used, where were part of ‘large group treatment’ of Reyes-Contreras *et al*. (2023). Breeding groups consisted of a dominant pair and eight helpers of varying sizes (4 large, 2 medium and 2 small helpers; expect one group, 52HH, which had 6 large helper and one medium and large helper respectively) and sexes. The standard length (SL) of breeding individuals ranged from 5.3 cm to 7.3 cm. Helpers were categorized into three size classes based on body length: small helpers (SL = 1.5–2.5 cm), medium helpers (SL = 2.6–3.5 cm), and large helpers (SL > 3.5 cm). Helpers were unrelated to the breeder pair and included immature and mature individuals of both sexes, reflecting natural group compositions. Some tanks contained small offspring recently produced by the breeding pair, which was noted during the experiments. Groups were housed in 300-L sections (97 × 65 × 50 cm) of 400-L tanks. The tanks were furnished with a 2-cm sand substrate, one half flowerpot per individual as shelters and potential breeding sites, mimicking natural conditions in which individuals maintain and defend personal shelters. Additional elevated hiding places were mounted near the water surface (semi-transparent plastic bottles),

### Task presentations

Two helping behaviours were elicited in group members by presenting them experimentally with helping challenges. These presentations were part of another study by Reyes-Contreras *et al*., (2023), who presented and video-recorded the helping tasks either simultaneously or successively in randomized order. For this study only recordings of simultaneous presentations were analysed. The two elicited helping behaviours included (1) territory maintenance by digging away sand and (2) defence against the egg predator *Telmatochromis vittatus.* Both helping tasks were presented near the shelter occupied by the dominant female (which is the most likely place of egg deposition in case the female would spawn). (1) In natural conditions, movement of sand by underwater waves can block shelter entrances. Fish can ‘dig’ out sand from the breeding shelters with their mouth or by vibrating their caudal fins and body, which is an important duty of brood care helpers in *N. pulcher*. To elicit digging behaviour, we covered the entrance of the breeding shelter up to 2/3^rds^ with sand. (2) To elicit defence behaviour against egg predators, we placed an individual *T. vittatus* in a clear Plexiglas beaker (radius = 5.25 cm; height = 15.5 cm, with perforations on the lid and on one side) near the entrance of the breeding shelter. In this set-up, defence against *T. vittatus* consisted of aggressive behaviours towards the Plexiglas beaker. In nature, eggs and larvae are vulnerable to both movement of sand into shelters and egg predation by *T. vittatus*. Groups were acclimated to the presence of the beaker for at least one week before trials started. Each trial lasted 20 min (5 min acclimatisation + 15 min behavioural recording) and were repeated three times per group to determine if the behaviours were consistent. Repeats were separated by at least one week. All trials were video-recorded using a hand-held camera (SONY® Handycam®, Model HDR-CX405) mounted on a tripod while the observer stayed outside of the aquarium room.

### Video analysis

The 15-min videos were analysed using the behavioural event-logging software BORIS (Friard & Gamba, 2016) adhering to an ethogram developed for this species (Reyes-Contreras *et al*., 2019). Each video was analysed repeatedly for five times, recording the size classes which performed a behaviour and the frequencies of behaviours. We recorded (1) the frequency of defence by group members (overt aggression exhibited as bites directed towards the beaker involving physical contact), (2) frequency of digging by group members (movement of sand from one location to another with the mouth), (3) overt aggression of the breeding male to the breeding female and subordinate helpers, (4) overt aggression of the breeding female to subordinate helpers, and (5) the activity level of *T. vittatus.* Activity level was recorded every 60 sec for 5 sec and was coded as 1 (movement occurred) or 0 (fish was inactive), resulting in an overall activity score ranging from 0 to 15 during a 15-min video.

In the behavioural analyses, we did not correct for the number of individuals in each size class because we could not track different individuals within size classes – we could only distinguish them by size. Yet, importantly, there were no instances were fish of the same size class performed the same or both tasks at the same time or in quick succession. Therefore, all reported task transitions within trials are within the same size class and same individual. However, while the task specialisation metrics, based on cumulative task performance, did not indicate specialisation between size classes (see ‘Results’), this does not entirely preclude the existence of specialists within individual size classes.

### Statistical analyses

All analyses were conducted in R (R 4.4.3, R Core Team, 2021). We first visualized all the tasks performed by the respective groups by plotting frequencies of digging and defence across five categories of fish, breeding female, breeding male and the three size classes of helpers. In all analyses, we did not account for when the group sizes changed due to evictions (13 cases across 33 trials), as these were remedied quickly by introducing a replacement individual and making sure that there was always individuals of all size-classes present.

#### 1. Task performance across size-classes

To analyse whether different size classes varied in their performance of digging or defence, we fitted generalized linear mixed models from the ‘lme4’ package in R (Bates *et al*., 2015). We fitted two Poisson Generalized Linear Mixed-Effects Models (glmm) with the counts of digging and defence counts as the response variables, size class as fixed effect and group identity as random effect, since groups were repeatedly tested three times. We analysed whether models were overdispersed using the ‘Dharma’ package (Florian et al. 2017). Models were not overdispersed and we report the estimates in log-scale, with their 95% confidence intervals, considering an effect to be significantly different from random when the 95% CI does not overlap with 0. Furthermore, we conducted pairwise Tukey’s post-hoc contrasts between the different size classes using the package ‘emmeans’ (Lenth 2022). To check if within the groups, the size classes were consistent in task performance over the three repeats, we computed the Intra Class Correlations (ICC) per group and performed an F-test with the null hypothesis that there is zero consistency, using the ‘irr’ package in R (Gamer *et al*., 2019).

Further, to understand whether different size classes were specializing on tasks, which can occur independently of whether or not there was division of labour, we computed the Proportional Similarity Index (PSi). PSi is widely used to analyse the diet specialisation or niche specialisation of individuals with respect to the population (cf. Marklund et al. 2018; Pagani-Núñez et al. 2015) and is also used to determine individual task specialization (cf. in common wasps, *Vespula vulgaris*; Santoro, 2015). We first computed the proportion of digging of each size class within a trial by dividing the digging performed by a size class by the total digging performed in the trial. Similarly, we computed the proportion of defence by each size class within a trial. Next, we calculated the average proportion of digging and defence across all trials as

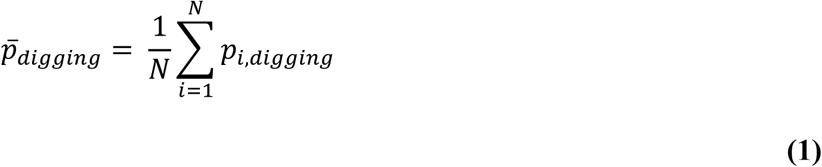

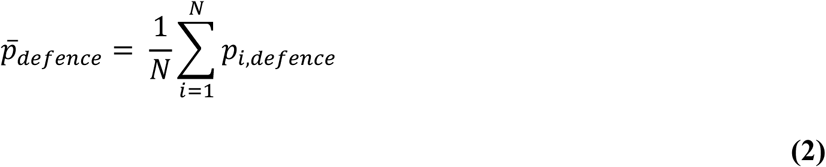

Finally, we computed the proportional similarity index for each size class *i* in a trial as

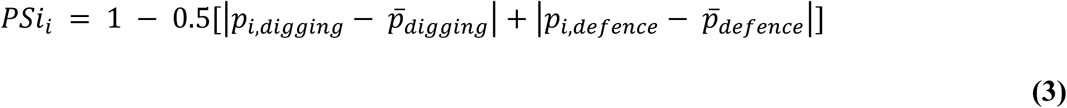

#### 2. Division of labour across size-classes

In the next step, we computed Normalised Mutual Entropy (NME) as a measure of DoL in our study groups (Gorelick et al 2004). NME is a useful index of DoL used extensively to compare division of labour in a standardized way across groups of different sizes and tasks distributed among them. We first calculated the marginal entropies, which measure the spread of tasks and workloads.

Marginal entropy is expressed as

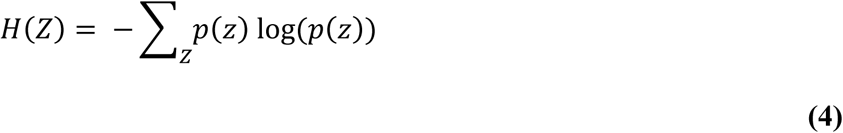

Marginal entropy across size classes is given by

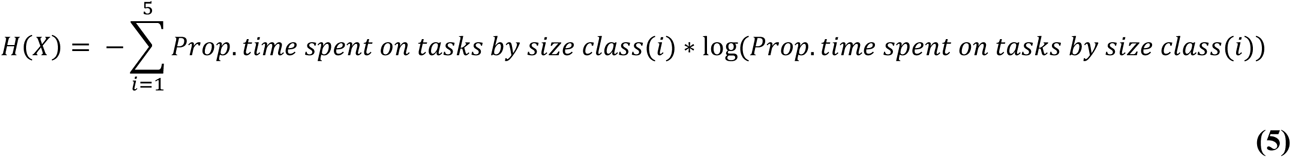

Marginal entropy across tasks is given by

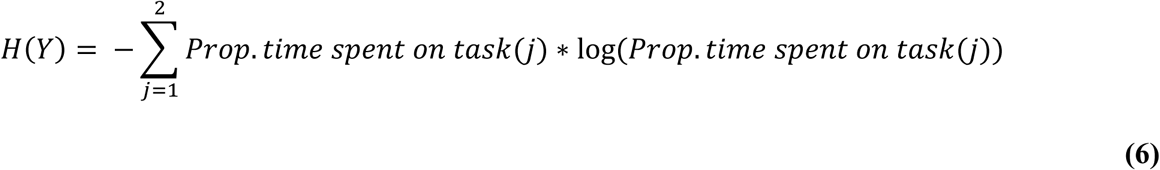

*H(X)* is the marginal entropy across size classes and describes how the total task effort is distributed across them. *H(X)* = 0 indicates that all tasks were performed by a single size class and high values indicate that the efforts were shared evenly across all size classes. Similarly, *H(Y)* is the marginal task entropy, which describes how workload is distributed across the different tasks. *H(Y)* = 0 indicates that all the efforts were focused on one task and high values indicate that the workload was evenly distributed across tasks (here, across digging and defence).

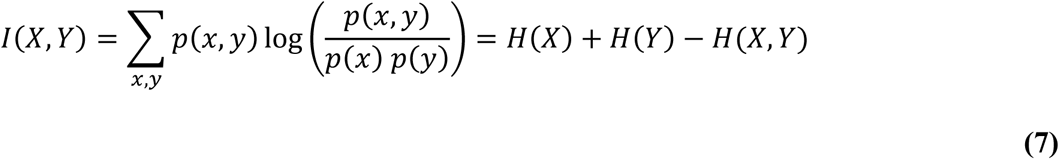

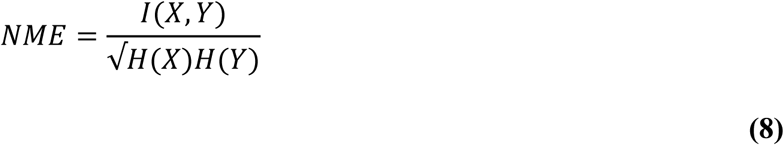

Mutual entropy *I(X,Y)* quantifies the amount of information that can be obtained regarding the tasks by considering the individual size classes, and conversely, the amount of information regarding the size classes that can be inferred from the tasks. Finally, symmetric division of labour, measured as Normalised Mutual Entropy (NME) normalises the mutual entropy by the geometric mean of the entropies, giving equal weight to both task and size class distribution and bounds the values between 0 and 1. A value of 0 indicates no division of labour and 1 indicates perfect division of labour across the size classes. Some groups attained DoL values of ∞ when either *H(X)* = 0 (one size class does all the work, or no work was done) or *H(Y)* = 0 (the total work done was either zero or individuals performed only one task in that round) leading to a division by zero. These groups were assigned DoL values of NA (not available).

#### 3. Patterns of task transitions within trials

In order to understand how the performance of different tasks is organized among group members, we analysed the transitions between tasks by the same or different size-classes. For this we used made use of the high time-resolution of our data, where we had recorded the exact times for each helping event within each trial. Individual size-classes were highly consistent within a trial in performing tasks and hardly switched between tasks (15/165 instances, 11 groups x 3 rounds x 5 classes) with the breeding male and female responsible for 12 of these switches. Therefore, we focused on how group members in general transitioned between tasks, without distinguishing further between size classes. We examined whether group members transitioned to the same task as performed by others s (‘Task contagion’) or to the other task (‘Spontaneous DoL’).

We had many instances of ‘idle’ individuals and rather few individual transitions from one behaviour to the next, preventing us from using probabilistic models to predict the next state of the system or the next event. Instead, we used lag-sequence analysis to analyse task transitions between individuals by counting the instances of the four possible transitions between the two tasks Digging and Defence, which are, Dig → Dig, Dig → Defence, Defence → Dig and Defence → Defence. From these counts, we tested whether transitions are more likely between same tasks (Dig → Dig and Defence → Defence) or a different task (Dig → Defence and Defence → Dig), using a binomial test. The detailed procedure and analysis are given below.

A visual inspection of task performance of digging and defence by size classes revealed that tasks were generally performed in ‘bouts’, wherein the same tasks are done in quick succession by the same individual (belonging to the same size class), with periods of longer breaks in-between. Hence, we first cumulated single ‘events’ of task performance into bouts per individuals per task type. To define the duration of a bout objectively, we employed a hazard rate approach. This method identifies the likelihood of an individual continuing the same task over time and estimates mean bout duration based on survival properties of behaviour.

##### Calculation of mean bout duration

We computed the mean bout durations for digging and defence events separately. As an example, here we compute the mean bout duration of digging events as follows: For each individual, we calculated the time interval (Δt) between consecutive events of task occurrence of the same behaviour (Dig→Dig). We partitioned these intervals into 1-second duration bins, to allow us to calculate bout duration in seconds. We then tallied how often transitions from Digging occurred within each duration bin: Dig→Dig, Dig→Def. Using this, we computed the hazard rate or likelihood for continuing the same task, Digging, as

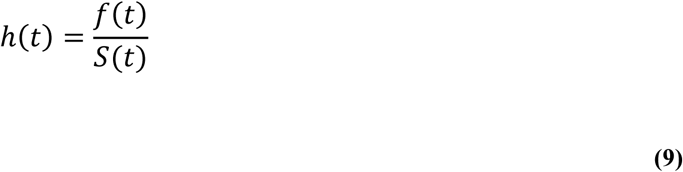

where *f(t)* is the number of observed same-task transitions (Dig→Dig) in bin *t* and *S(t)* is the cumulative number of individuals “at risk of ending the bout” - that is, individuals still performing that task (Dig) or which have switched to another task (Dig→Def) at or beyond time t. We then plotted the hazard rates across time separately for digging and defence (Figure S1). For both digging and defence, visual inspection of the hazard plots indicates that the hazard rates of task performance reach persistently low values after ∼10 s. That is, until 10 s after performing a task, individuals have a high likelihood of performing the same task and therefore can be seen as performing the task in a bout. Therefore, events of the same task performed by an individual within a ≤10 s interval after the previous event are considered to belong to the same bout. If an individual performs the same task >10 s after its previous event, or it switches to a different task this is considered a different bout (Figure S1 & S2)

##### Follow events

To characterize the succession and coordination of task performance, we then identified between-individual ‘follow’ events. We considered that an individual ‘followed’ another one when it performed a task within 5 s after the initiator’s bout ended. These ‘follow events’ were categorized by task pair (e.g., Dig→Dig, Dig→Def, Def→Dig, Def→Def). We arbitrarily assigned a 5-s window as a reasonable timeframe to detect responsive behaviour from the ‘following’ individual (Figure S2). To make sure our results are not an artefact of this window, we performed a sensitivity analysis by choosing different windows (3, 5, 10, 15 and 20 s). We show that the results are similar over different window lengths (Figure S3).

## Results

### Overall task performance

Across all trials, individuals of all size classes performed both tasks at least once, indicating that they are all capable of performing the tasks. Within trials, size classes were consistent in their task performance and rarely switched between tasks. Task switching by the same size-class within a trial occurred only in 15 out of 165 cases (11 groups x 3 rounds x 5 size classes), with breeding pair male and female responsible for 12/15 cases. In 7 out of 33 trials, none of the group members performed any of the tasks. Digging and defence tasks were performed on an average 5.01 (±16.07 SD) and 3.50 (± 12.94 SD) times across all trials.

### Size-based task performance

Overall, dominant breeding females performed the most tasks across all trials (Figure 1). Breeding females dug sand more often and breeding males defended more often against the egg predator when compared across all size classes (pairwise contrasts in log-scale, Table 2 and Figure 1). Task performance across size classes in the groups was not consistent over the three repeats (digging: mean ICC = 0.18, SD = 0.28; defence: mean ICC = 0.10, SD = 0.23, Table 3), except in three groups, which showed moderate to high consistency.

**Figure 1.**
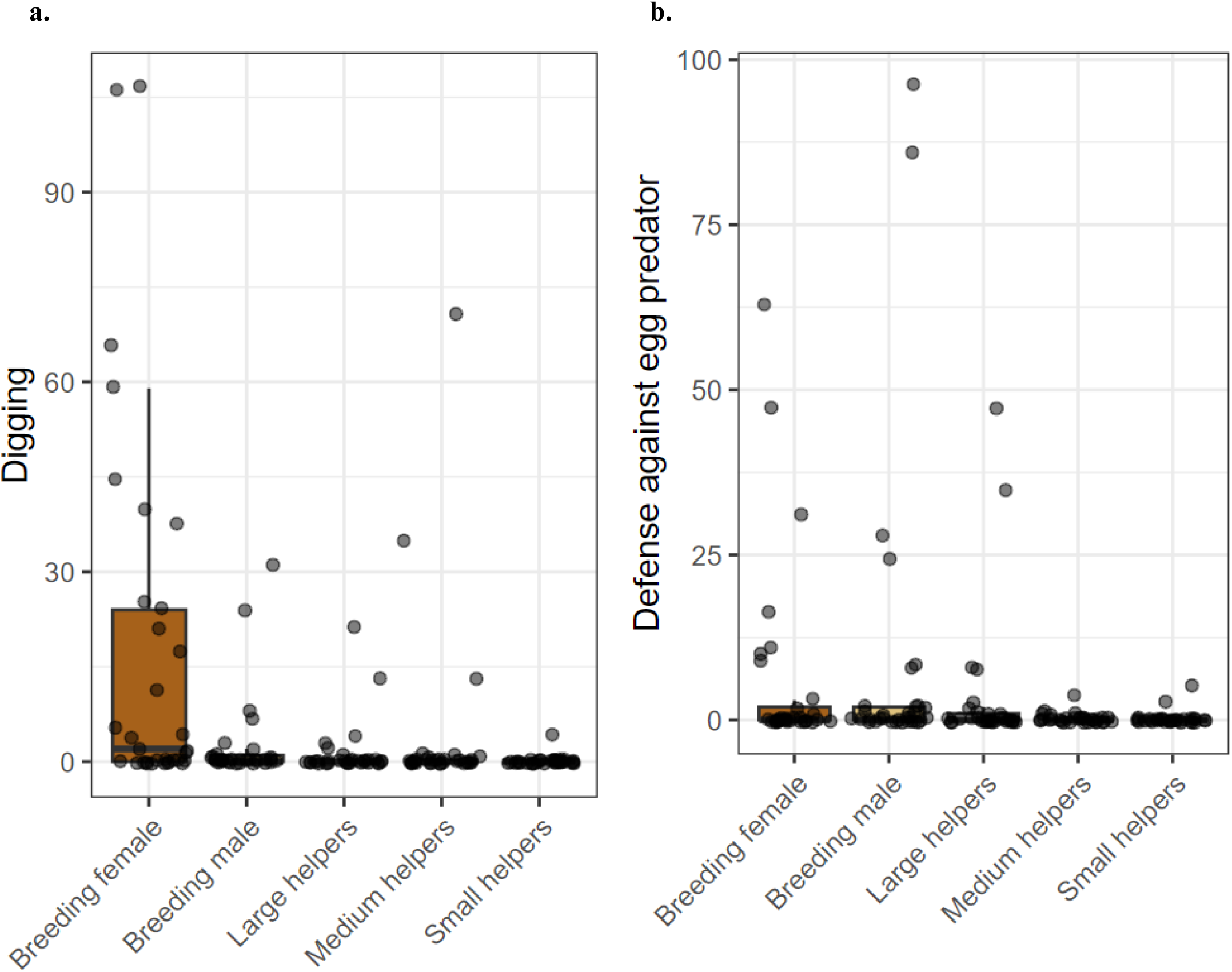
Task performance across size classes across all groups in the cooperatively breeding cichlid *N.pulcher* (N=11 groups). Boxplots show the total number of digging events (a) and defence responses (b) towards the egg predator (*T.vittatus*) performed by different size classes, pooled across all groups and rounds. Dots represent individual-level data, and boxplots indicate median and interquartile range

**Table 1.**
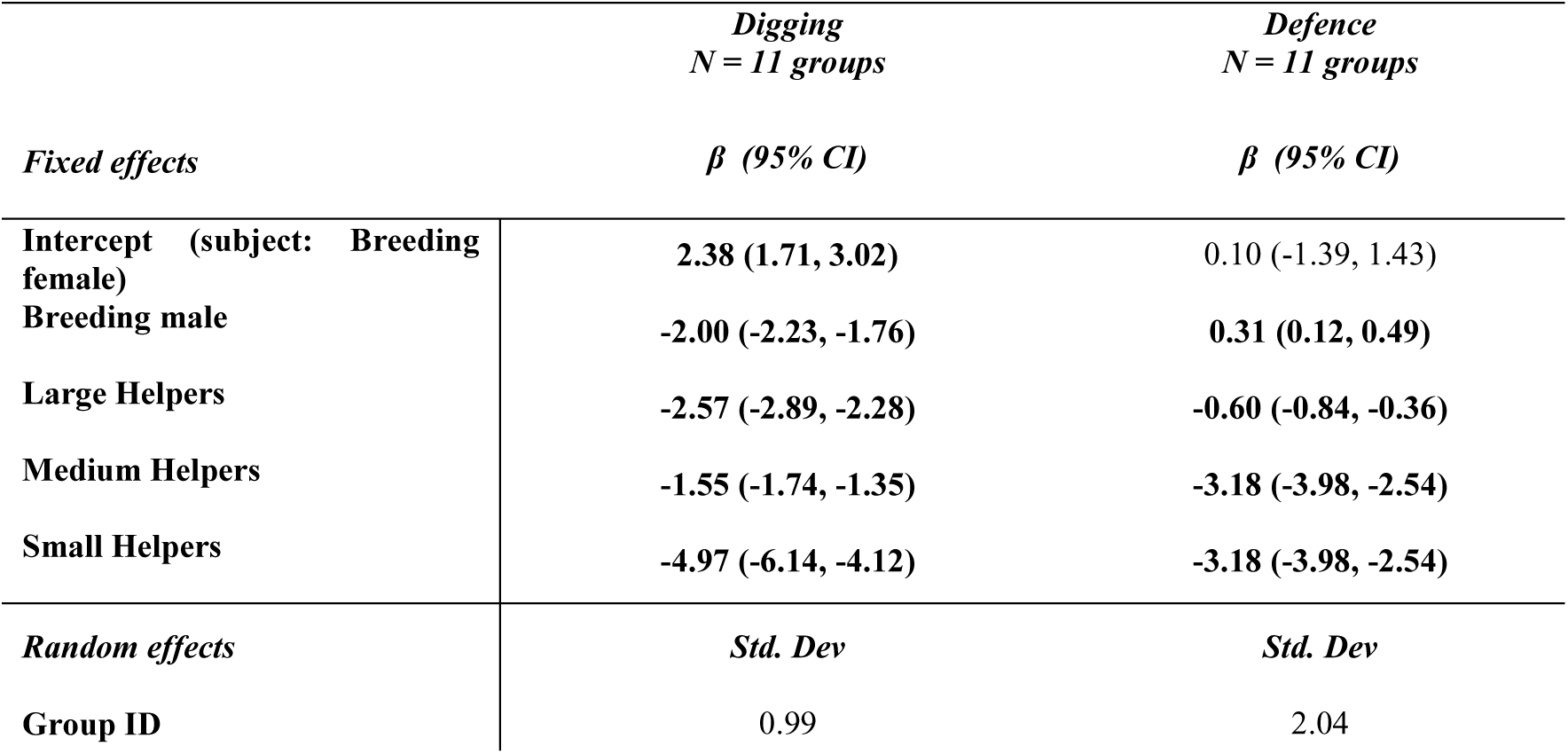
Effects of size class on task performance. Summary of generalised linear mixed model (glmm) results showing the effects of size class on two cooperative tasks: digging and defence against an egg predator. β values are the estimates in log-scale and 95% CI are indicated within the brackets. Significant estimates, where the 95% CI does not overlap zero, are set in bold

**Table 2.**
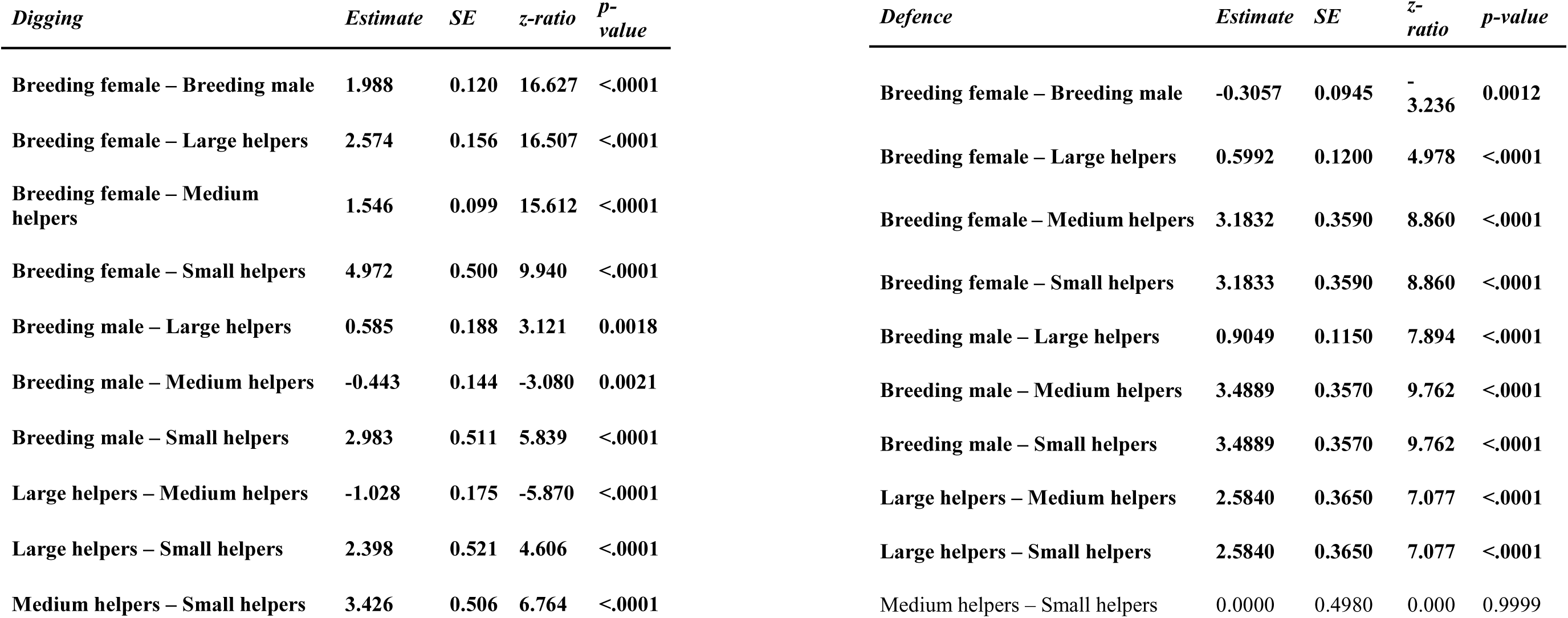
Post-hoc pairwise comparisons of task performance across size. Results of Tukey-adjusted pairwise comparisons using estimated marginal means (emmeans) following generalized linear mixed models from Table 1 for digging and defence against an egg predator. Significant contrasts (p < 0.05) are indicated in bold

**Table 3.** Intraclass correlation (ICC) of digging and defence behaviour across size classes in three repeats per groups. ‘Significant’ ICC values > 0.75 are considered as high consistency, values between 0.4 and 0.75 as moderate and values between 0.1 and 0.4 as low consistency (in bold). Non-significant ICC values are considered to have very low or no consistency. Negative ICC values indicate very large variation

| Digging |  |  |  | Defence |  |  |  |
| --- | --- | --- | --- | --- | --- | --- | --- |
| Group | ICC | p-value | Interpretation | Group | ICC | p-value | Interpretation |
| 43AA | ~0 | 0.461 | No consistency | 43AA | ~0 |  | No data |
| 51BB | 0.008 | 0.450 | No consistency | 51BB | ~0 | 0.461 | No consistency |
| **52HH** | **0.496** | **0.047** | **Moderate consistency** | 52HH | ~0 | 0.461 | No consistency |
| **55X** | **0.763** | **0.003** | **High consistency** | 55X | 0.315 | 0.138 | No consistency |
| 57JJ | 0.058 | 0.387 | No consistency | 57JJ | 0.362 | 0.108 | No consistency |
| 59U | ~0 | 0.461 | No consistency | 59U | 0.242 | 0.194 | No consistency |
| 62LL | ~0 | 0.461 | No consistency | **62LL** | **0.484** | **0.051** | **Moderate consistency** |
| 68PP | 0.305 | 0.145 | No consistency | 68PP | -0.166 | 0.69 | Very large variation |
| 69QQ | 0.016 | 0.441 | No consistency | 69QQ | ~0 | 0.461 | No consistency |
| **70RR** | **0.454** | **0.062** | **Moderate consistency** | 70RR | -0.208 | 0.749 | Very large variation) |
| 71SS | -0.076 | 0.564 | Very large variation | 71SS | ~0 | 0.461 | No consistency |

### Division of labour and task specialization across groups

Division of labour across size classes, calculated as the NME, varied greatly among groups, ranging from very high values near 1 indicating strong division of labour (group 52HH) to nearly zero (group 62PP; Figure 2). The large variation across groups with similar group sizes and compositions of helpers suggests that labour is flexibly allocated within *N. pulcher* groups. The PSi values, estimating task specialization, indicated that the different size classes were generalists with Psi values close to 1, except for breeding females and breeding males, which showed moderate specialisation in digging and defence respectively (median PSi for breeding females: 0.68 and breeding male: 0.58, Figure 3).

**Figure 2.**
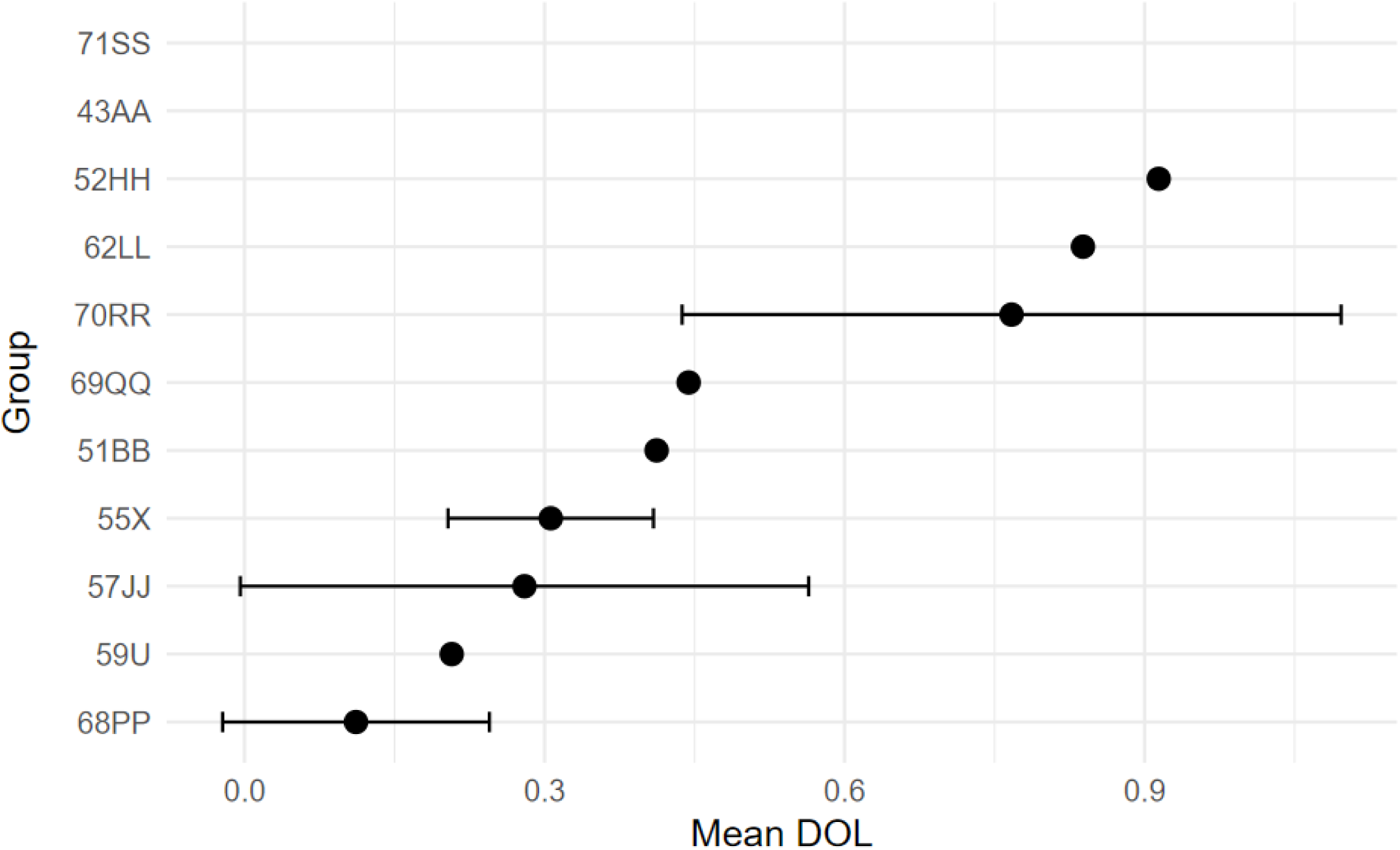
Normalised Mutual Entropy (NME) as an index of division of labour. The dots show the mean DoL of a group across the three trials and the whiskers indicate the standard deviation. For each group, DOL values were calculated per round. The DOL values for 71SS and 43AA are NA, which indicates that for each group and round, the total behaviour events (Digging + Attack) were either zero or individuals performed only one task across rounds

**Figure 3.**
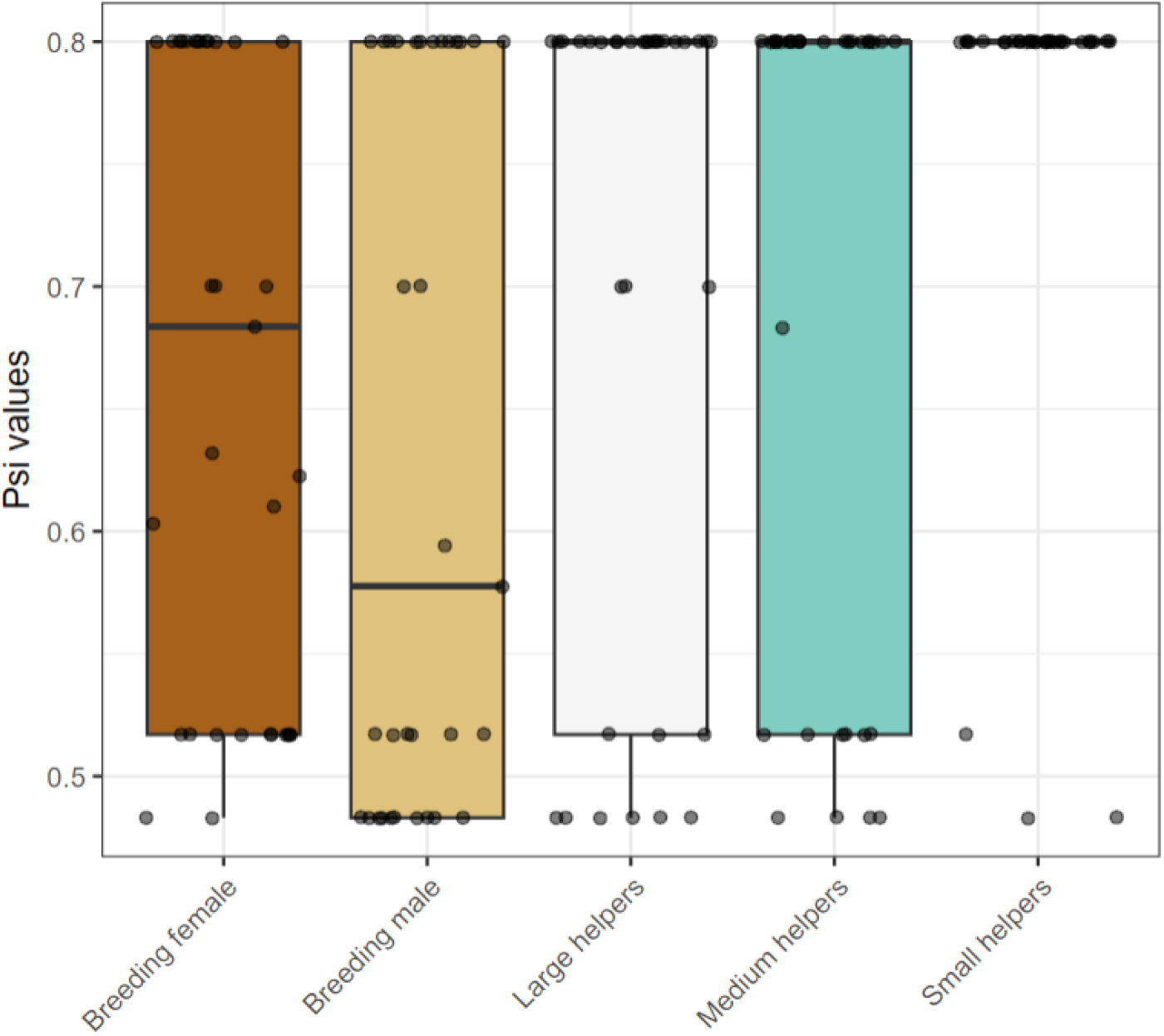
Proportional Similarity Index (PSi) values across social classes under two task-weighting conditions. PSi values close to 1 indicate behaviour closely matching the group average or low specialization and values closer to 0 indicate divergence from the group average or high specialization within the size class. Boxplots represent the distribution of individual PSi values within each size class; dots indicate individual data points

### Patterns of task performance within trials

Individual size classes were more likely to ‘follow’ another size class in performing the same task (’task contagion’, probability 0.82) rather than spontaneously divide their labour and take up the other task (probability = 0.82; binomial test, p < 0.001), Figure 4).

**Figure 4.**
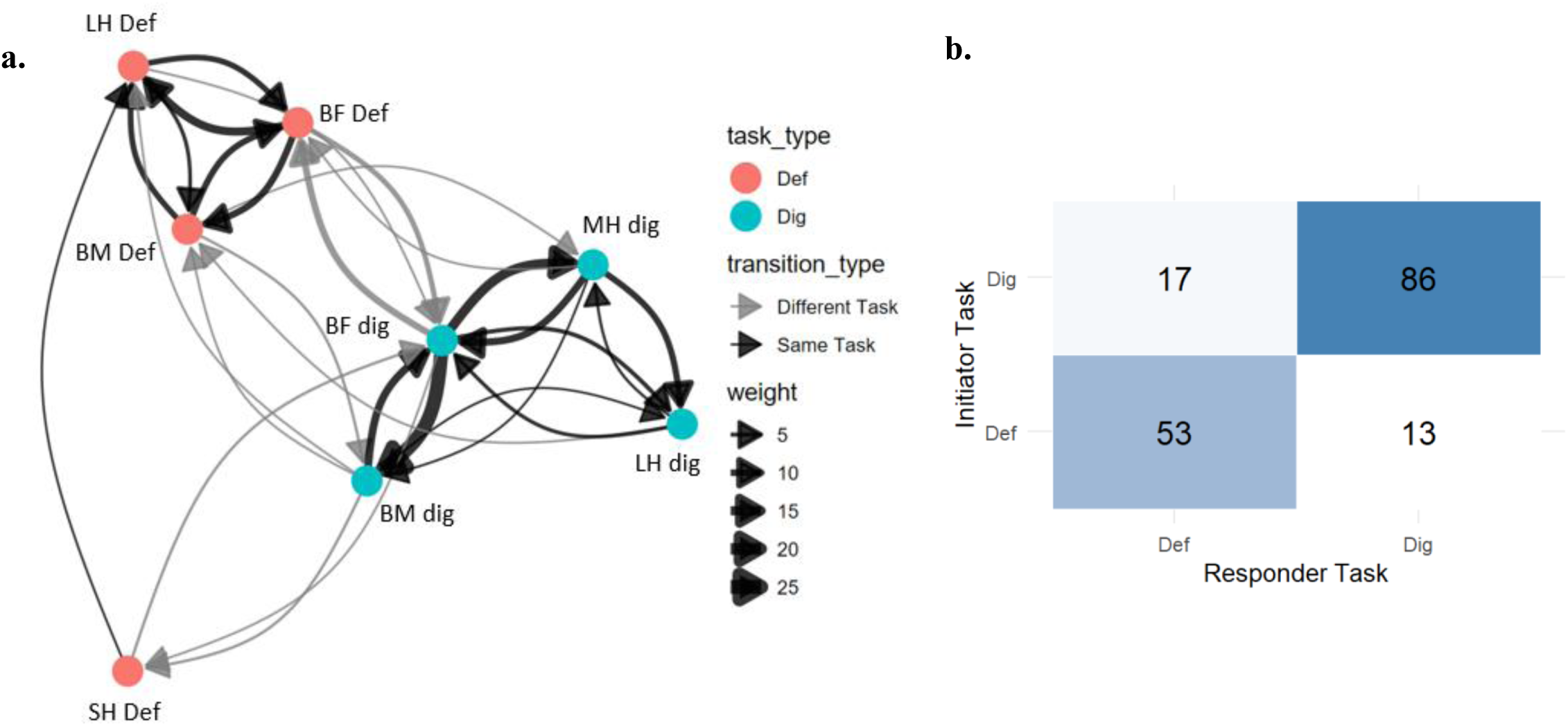
a. Across-size class transitions in performing digging and defence tasks. For each task bout performed by an initiator size class, we searched for subsequent behaviours by responder size classes within the same group and round that occurred within a 5-second following time window. Only pairs of behaviours from different size classes were included as individuals of a given size class tended to perform only one task type within a trial (169 follow events in total). Black arrows represent same task transitions and grey arrows represent transitions to a different task. Thickness of the arrow represents transition counts and the different colours represent the different tasks. b. Frequencies of observed transitions between same behaviours (Dig→Dig and Def→Def) and different behaivours (Dig→Def and Def→Dig). Note that ‘initiator’ and ‘responder’ are termed solely for simplicity of understanding and are defined based on which individual/class performed a task first and which performed subsequently. They do not indicate social interaction or intentional behaviour.

## Discussion

In this study, we find evidence that task allocation and division of labour is flexible across similar groups of *N. pulcher,* across trials (see example of different patterns in Figure S4 and a detailed overview in Figure S5). Groups mainly contained generalists, with the breeding female performing most of the tasks, with moderate specialisation in digging and the dominant breeding male defending marginally more than the dominant breeding female. Across multiple trials DoL was very low and variable. In some trials, entire groups failed to perform any cooperative tasks, underscoring high variability in task engagement.

We further uncover a potential mechanism of task performance and coordination of tasks as a group in this system. Individuals are more likely to follow each other in performing the same tasks, which fit the idea of a behavioural or ‘task contagion’. This phenomenon may represent a lower-level coordination rule, which, under certain conditions, can facilitate effective task organisation without complex communication.

### Lack of size/age based DoL

In our study, we observed significant inter-group variability in the division of labour (DoL) among groups of *N. pulcher*, despite identical demographic compositions (Figure S5). Some groups exhibited a high degree of task specialization, indicated by normalized mutual entropy values approaching 1, while others demonstrated little to no division of labour, with entropy values near 0. This disparity suggests that the emergence of DoL is not a straightforward outcome of group structure but may be influenced by contextual factors, motivational differences, or due to chance occurrences. This contrasts with previous field experiments indicated that *N. pucher* exhibited size polyethism, with larger individuals specializing in tasks such as digging and smaller helpers defending against egg predators (Bruintjes & Taborsky, 2011). Three potential explanations could account for this discrepancy: first, the stable lab environment may have reduced the perceived need for help, allowing dominants to perform most tasks themselves and leading to low frequencies of helping behaviours by group members in general; second, the specific nature of tasks may have influenced participation, with groups prioritizing either defense or digging based on immediate threats or priorities (even though the observed low engagement in tasks does not support a prioritisation of tasks); and third, individual variation within size classes may obscure size-based specialization, as behavioural tendencies shaped by early-life social conditions could influence whether individuals are more likely to help or submit (Le Vin *et al*., 2011; Fischer *et al*., 2017). Overall, this is in line with patterns observed in various cooperative breeders, wherein DoL is not a fixed outcome, but a context-dependent phenomenon influenced by environmental factors, social interactions and stochasticity (Smith & Riehl, 2022).

### Functional significance of task contagion

In this study, we identified patterns of task performance that suggest a ‘task contagion’, where an individual’s propensity to engage in a specific task, such as task A, increases when another group member is also engaged in task A. While the same pattern could arise if there is a group-wide prioritisation of a task, low overall engagement across both tasks in our system makes that explanation less likely.

Behavioural or task contagion and social contagion, which involves the matching of behavioural actions and emotional and motivational states, respectively, among group members (see reviews Firth, 2020 and Pérez-Manrique & Gomila, 2022), is one way to synchronize group activities. At the individual level, task contagion may confer several benefits. It can be efficient to follow the behaviour of another individual engaged in a task already rather than assessing the current need for tasks oneself (information efficiency). Task contagion also may keep individuals being engaged in the same task by social reinforcement, thereby reducing switching costs (Chittka & Muller, 2009). Further, in a structured social system such as cooperative breeders, task contagion by mirroring the dominant’s behaviour could signal cooperation and maintain social ties and potentially reduce the aggression towards subordinates (Sandars *et al*., 2024). However, there are costs associated with behavioural or social contagion, including the potential spread of maladaptive behaviours, false alarms, or inappropriate stress or fear among individuals (Carnevali *et al*., 2020; Gray & Webster, 2023). Despite these risks, the benefits can outweigh the costs, as task contagion allows for rapid local recruitment to essential tasks (for example, mobbing a predator) and enables groups to effectively regulate task performance and synchrony. Coordination can emerge through low-level contagion and social facilitation, without the need for centralized control or other top-down mechanisms. Thus, while there are potential disadvantages of contagion, the resulting enhancements in group coordination and efficiency underscore its adaptive significance in cooperative systems. Such behavioural contagion in fear and predator evasion responses has been observed in zebrafish, and is regulated highly-conserved oxytocin pathways and social decision-making networks (Fernandes Silva *et al*., 2019; Burbano Lombana *et al*., 2021; Akinrinade *et al*., 2023; DeAngelis & Hofmann, 2023; Kareklas & Oliveira, 2024), which may also underlie ‘task contagion’ in *N. pulcher*.

### Conclusions

Understanding the behavioural mechanisms underlying coordination in task performance within such a complex system remains an important area for research. We have uncovered one such mechanism – task contagion, which can promote effective task performance. Simple behavioural rules such as task contagion may serve as precursor for the evolution of more complex DoL. In the case of simultaneous task demands as in *N. pulcher*, even minor variations in responsiveness to specific tasks or random initiation of tasks among group members can enhance recruitment, ultimately leading to successful task completion. In systems with large groups or spatially dispersed tasks, centralised coordination may be inefficient (for example in collective movement and collective decision making; Attanasi *et al*., 2014; Pacher *et al*., 2025). Under such conditions, local contagion could recruit nearby helpers rapidly, leading to emergent patterns of specialisation with or without DoL over time. Furthermore, it is possible that localised task demands and task contagion can result in individuals gaining exposure to a particular task throughout their ontogeny (sometimes even before they are able to perform the tasks themselves), adding to their skill in performing this task later on (Taborsky, 2025a).

## Acknowledgements

We thank Maria Contreras-Reyes and Guo Yijun for creating the groups, maintaining them and collecting the video recordings. Evi Zwygart for animal caretaking and logistic support, and the fish team and the Hasli team for discussion.

## Ethical note

All experiments were conducted at the Ethological Station Hasli of the Institute of Ecology and Evolution of the University of Bern, Switzerland. The captive holding of the fish was carried out under licence no. BE 4/22 and experiments were conducted under licence no. BE 34/22 of the Veterinary Office of the Canton of Bern. All fish were monitored daily during feeding by noting normal feeding behaviour, posture, morphology and behaviour. Evicted fish, recognised by limited movement and increased aggression by conspecifics were identified during daily monitoring and immediately isolated. All fish were retained in our permanent breeding stock after the end of the study.

## Data availability

The data, along with the codes that were used to produce the figures and analyses in the manuscript will be submitted in a Zenodo repository.

## Use of AI

ChatGPT was used to simplify code in R and for overall improvement of grammar. All contents were written by the authors.

## Consent for publication

All authors have given their consent for publication

## Supplementary material

**Figure S1.**
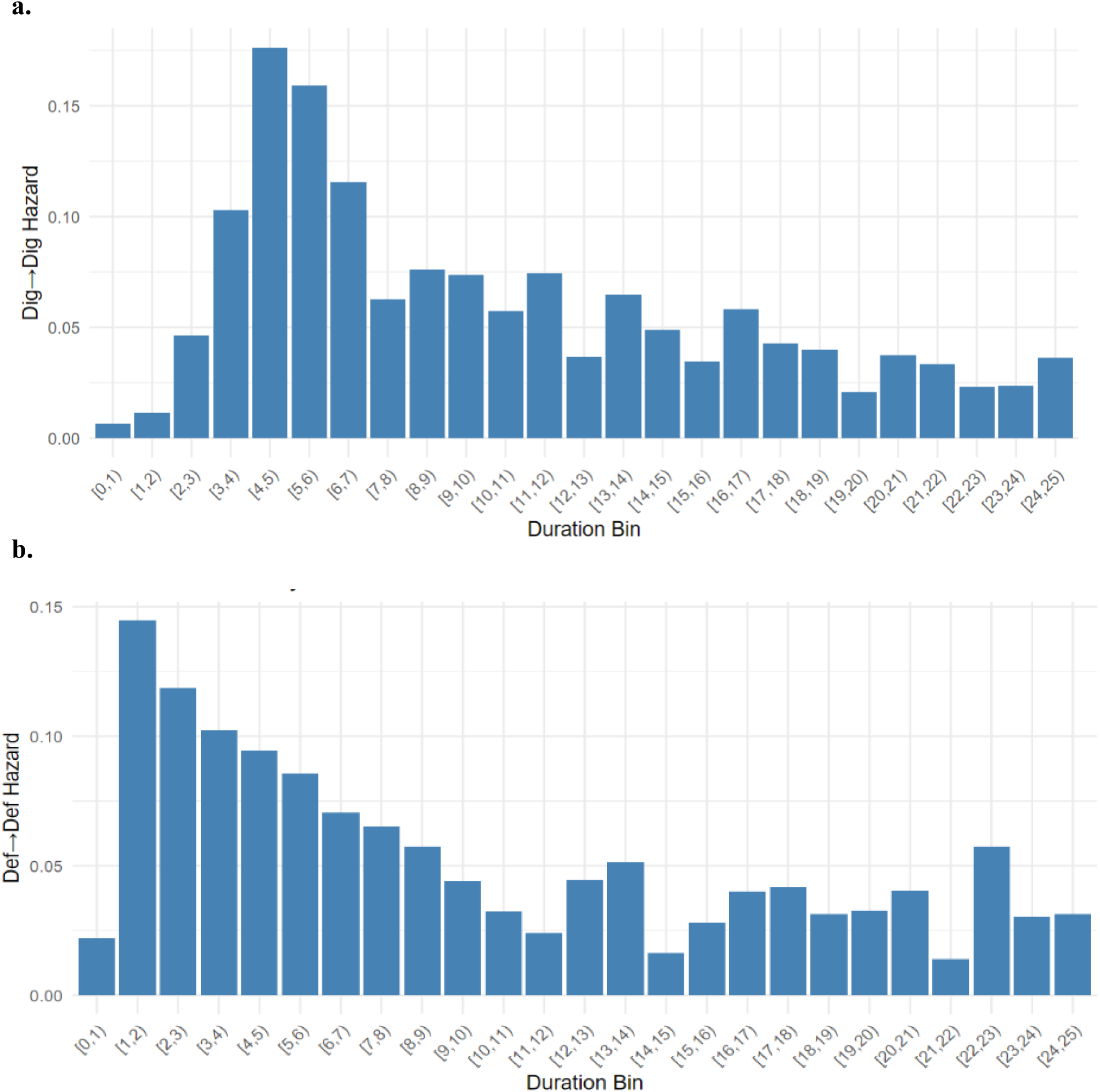
Hazard rates for specifying bouts for digging and defence events. The plots show the estimated hazard rates for behavioural persistence in (a) digging and (b) defence tasks. The hazard rate represents the conditional probability that individual transitions out of a given behaviour at time *t*, given that it has continued the behaviour up to that point. Both behaviours exhibit a steep decline in hazard rates within the first 10 seconds, after which the rates reach a relatively stable, lower baseline, which was used as a cut-off to determine bout length

**Figure S2.**
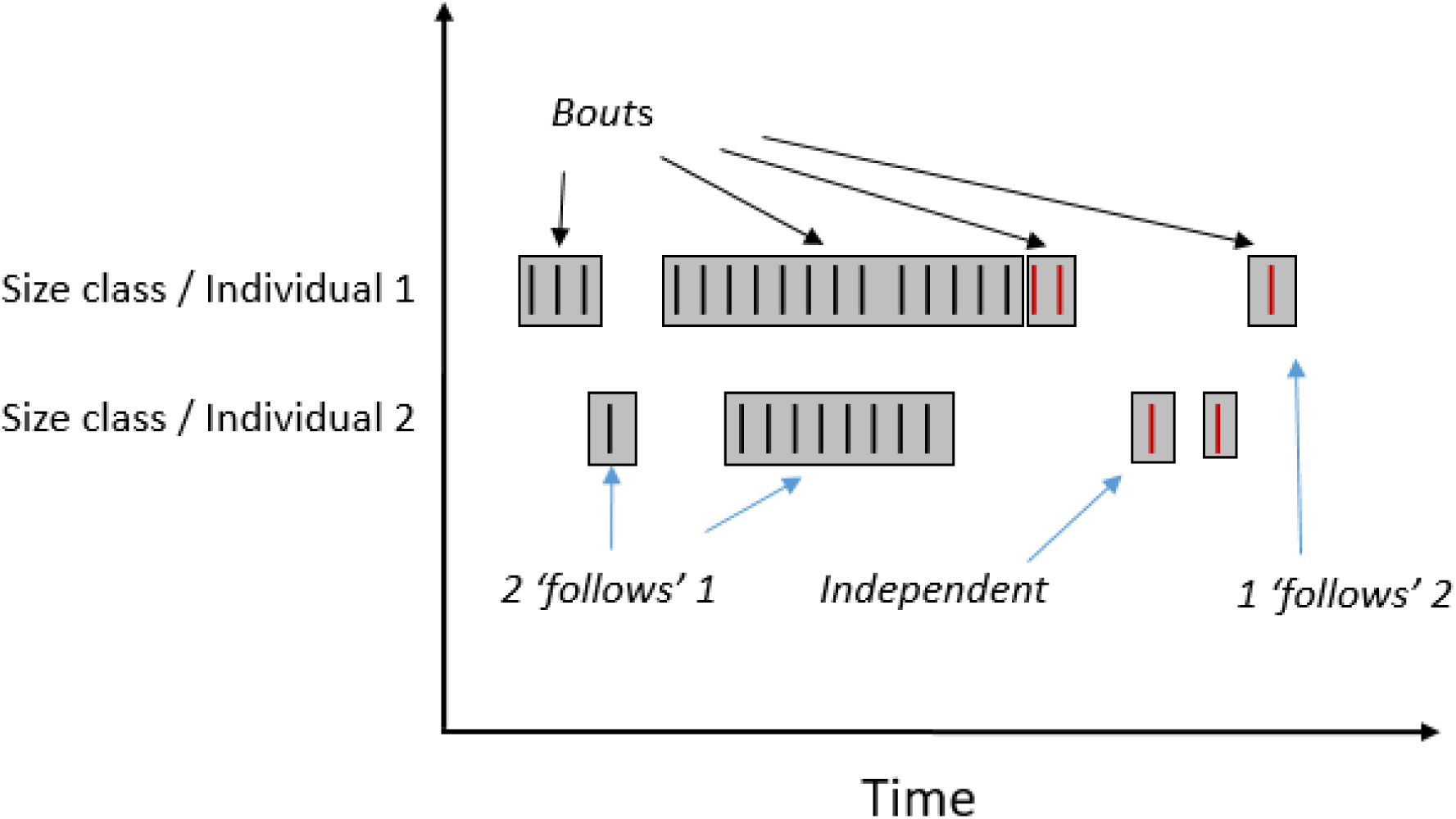
Classification of bouts and following events, illustrated with two hypothetical individuals. Bouts are classified as clusters of the same behaviour (e.g. black or red) performed by the same size class or individual within 10 seconds (cf. hazard rates in Fig. S1). A switch from one behaviour to another (e.g. black to red), even within 10 seconds, marks the start of a new bout. ‘Following’ events occur when a different individual or size class performs a behaviour within 5 seconds after the initiator’s action. Each bout includes an identified ‘initiator’ and ‘follower’, and these roles can vary across individuals or size classes within a single trial.

**Figure S3.**
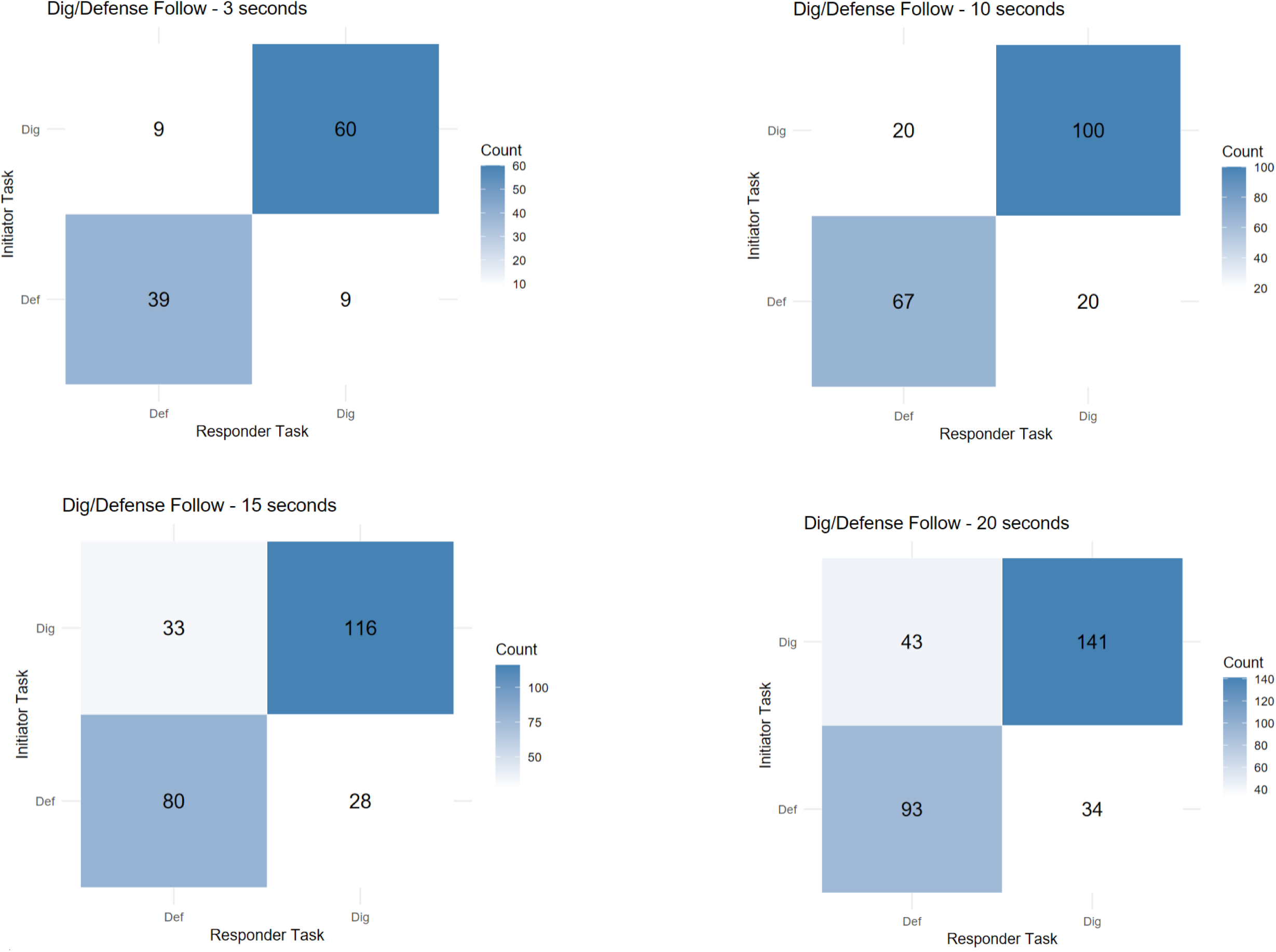
Results of the sensitivity analysis to assess the robustness of our operational definition of a “following event. We varyied the allowable time lag between an initiator’s action and the next action of a second individual. Specifically, we tested temporal thresholds of 3, 10, 15, and 20 seconds (5 seconds is the default in the main text), within which a subsequent behaviour by another individual or size class was considered as a follow

**Figure S4.**
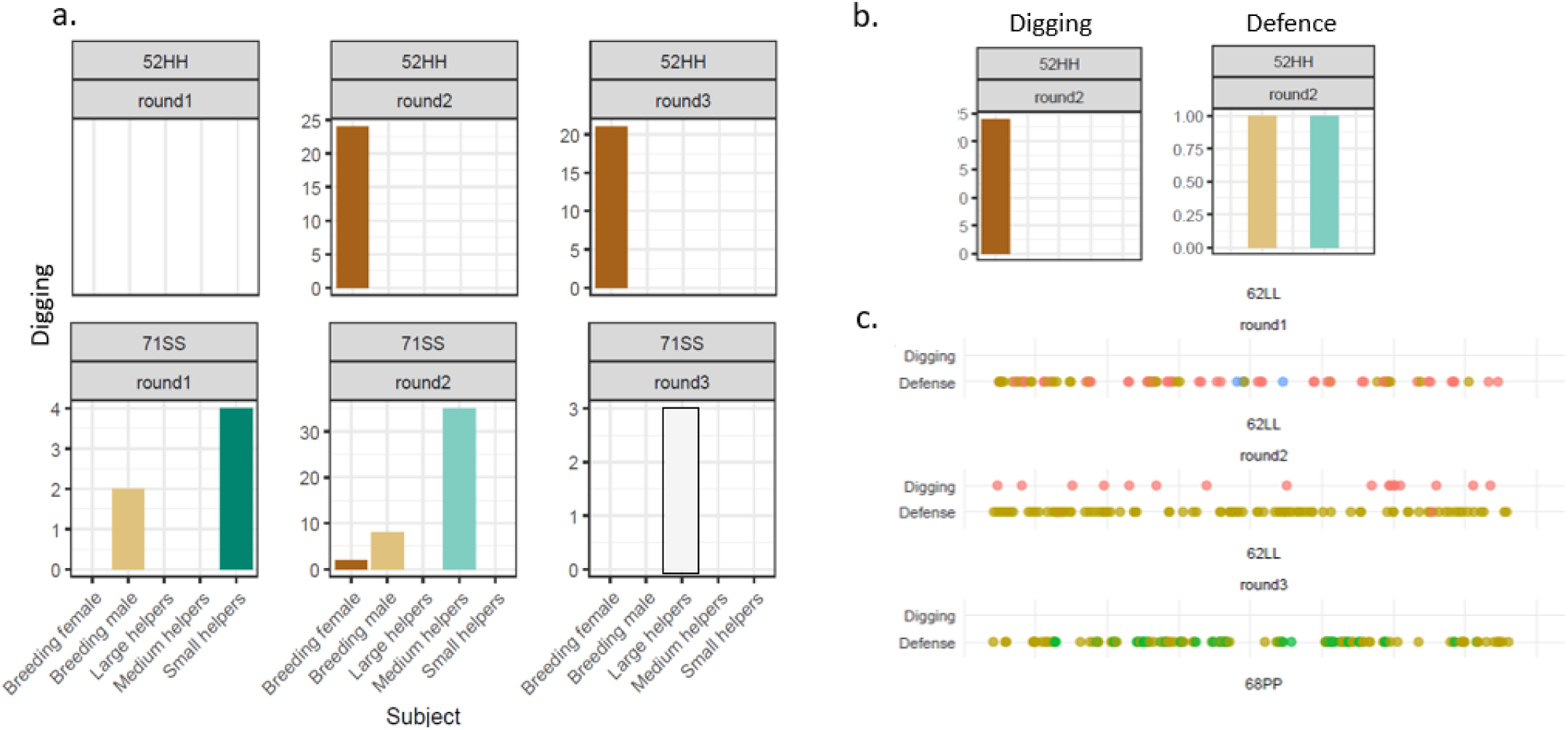
a. An example of a group (52HH) showing consistency in digging by size class across the last two rounds (there were no digging events in the first round) and of another group (71SS) with a lack of consistency across size classes. b. An example in which division of labour occurred in a group (52HH). c. An example of group 62LL, in which there was division of labour in round 2 but not in rounds 1 & 3 – plotted over time to visualise spontaneous DOL, different colours represent different helper classes.

**Figure S5.**
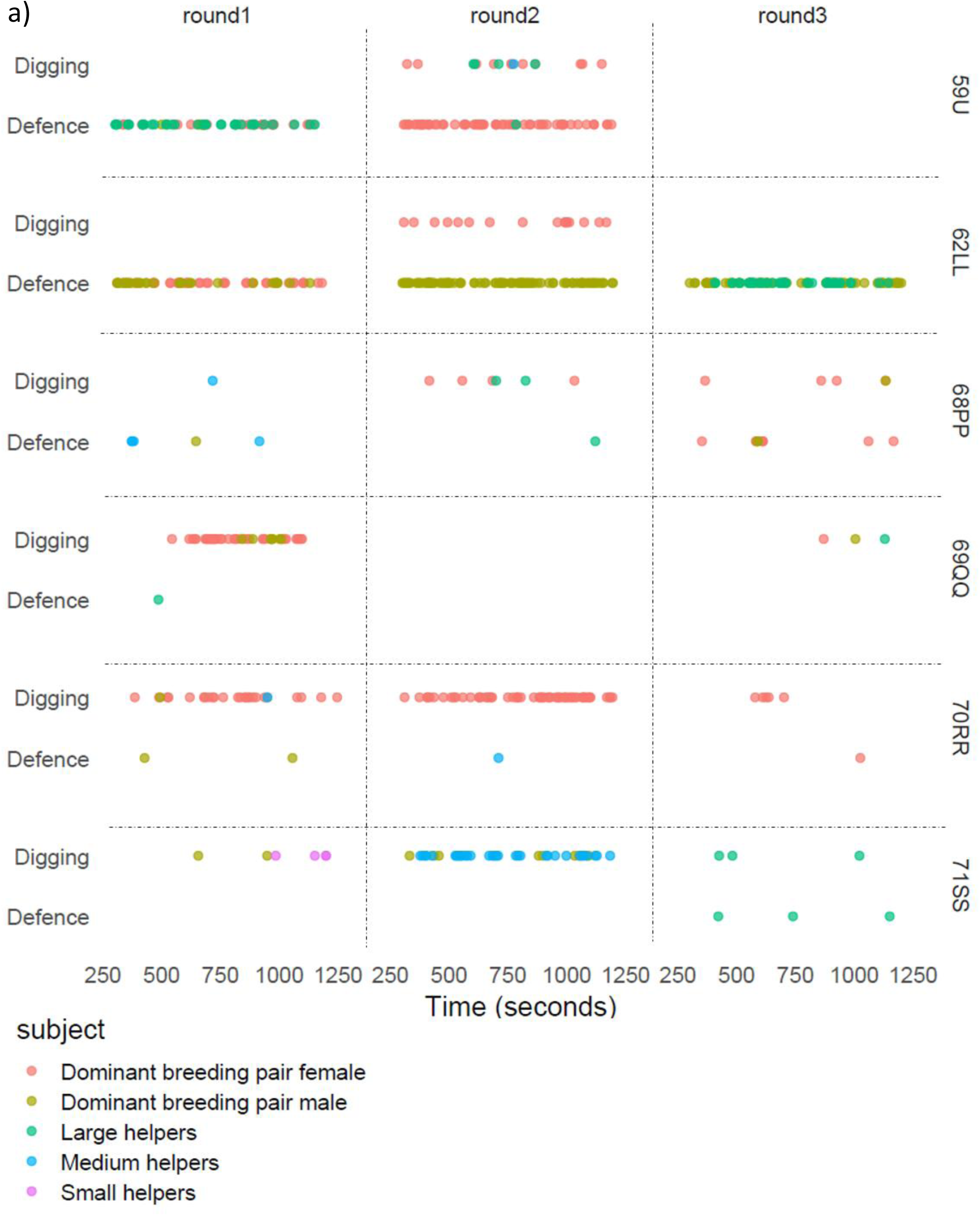
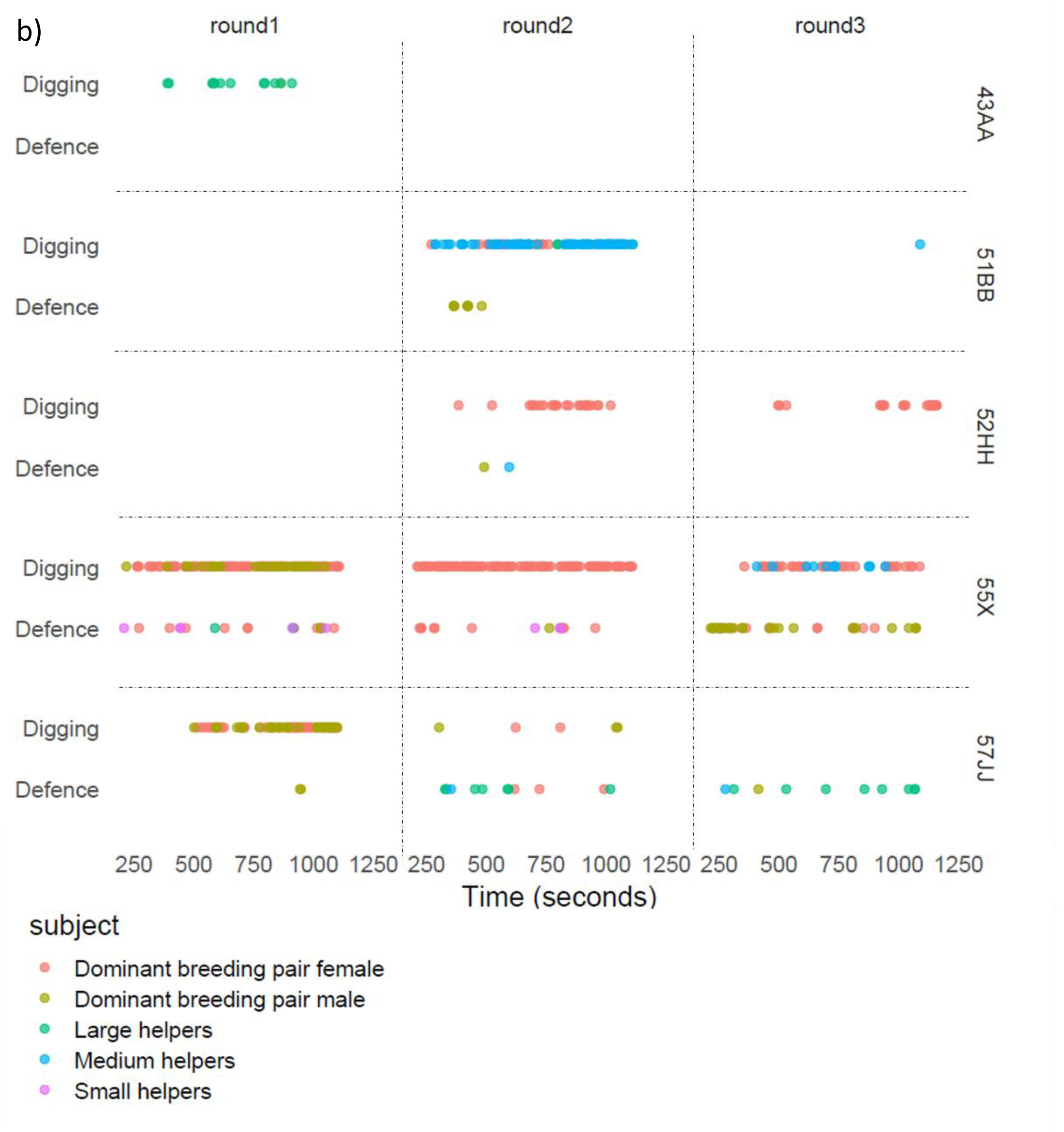
a&b) The two tasks performed over time across the rounds in different groups. This plot gives the overall high-resolution data collected within each trial for different groups. Each point represents an observation of task performed by an individual / size class given by the colour.

